# Super Quantum Neuroelectromagnetic Dynamics of Action and Percept Potentials

**DOI:** 10.1101/2025.02.24.639809

**Authors:** Jolly Kanjua

## Abstract

The spatiotemporal selectivity of neurons to sensory stimuli that underlie the center and surround receptive fields represents the anticommutative and, thus, symmetry that defines sensorimotor transformation. However, owing to the nonabelian nature of the irresponsiveness of neurons to sensory stimuli characterized by inexcitations, symmetry and, thus, transformation does not become apparent. Hence, a neuroscientific model that describes the precise mechanism of the nonabelian gauge group in the brain is unknown. Using quantum field theory, we show visuomotor transformation of natural images in which the superposition of opposite parallel cortical columns creates visual fields and annihilates motor fields. The creation and annihilation operators, magnetic and electric charged particles, are opposed inertia systems that act on each other by raising and lowering each other’s particles, leading to momentum–energy trade-offs, in our case, retinotopic gain and retinomotion loss, and vice versa. The magnetic charged particles *K*^−^, *Cl*^+^, *Na*^−^, intrinsic to a magnetic neuron, nabla, are predicted to exist opposite parallel to the electrically charged particles *Na*^+^, *Cl*^−^, *K*^+^ of the electrical pyramidal neuron in the neocortex. The implication is that the physical neocortex or electric brain is in concert with a nonphysical complex neocortex or magnetic brain that is hardly detectable but much alive and proactive.

## I. INTRODUCTION

Neurons with insensitive foveal regions and directionally sensitive extrafoveal areas were first reported in an early study of the waking monkey posterior parietal cortex^1^. These neurons, i.e., visual neurons, were dubbed light-sensitive neurons because the experimenters used light as stimuli (also images) to elucidate the functional properties of the neurons. These neurons were found to have the peculiar properties of foveal sparing: a rare response to fixation target stimuli even though the same stimuli, when delivered close to the fixation point, evoke a vigorous response. This finding indicates that stimulus properties such as the direction and velocity of motion at the extrafoveal areas are sensed, whereas stimuli at the center region are not. Such foveal sparing and opponent vector organization occur in the posterior half of the posterior parietal cortex, the inferior parietal lobe (IPL). In most of these cases, neurons are excited to the movement direction and velocity in the opposite direction of the fixation point and are inexcited to the fixation stimulus target. In contrast, neurons with sensitive center regions and insensitive extracenter areas have also been reported in separate studies of the monkey and cat visual cortex^2,3^. These neurons also have the properties of endstopping: a vigorous response to a particular length cutting off extras that might remain or extend beyond the center region. The stimuli used here were also light.

The outstanding question that remains to be answered is whether the excitability of one system influences the inexcitability of another system, linking the retina and the parietal lobe or the visual cortex and the parietal lobe or the motor cortex and the visual cortex or the retina and the motor cortex or all the above-listed cortices and thalamic nuclei to each other at once. In other words, the excitability and inexcitability between separate cortices cooccur, as excitability and inhibition cooccur within each cortex, performing linear transformation from retinotopic coordinates to generalized momenta or momentum space (previously believed to be spatiotopic coordinates). Simply, does foveal sparing result from endstopping causing the opponent vector effect, or does endstopping result from foveal sparing causing the extraclassical effect? Thus, straightforward interplay, a causal relationship of foveal sparing and opponent vector effects, endstopping and extrablassical movement, or foveal sparing and endstopping, linking different cortical areas, including visual and motor regions, has been difficult to elucidate.

Here, we present simulations suggesting that foveal sparing and endstopping effects may result directly from the superquantum neuroelectromagnetic dynamics of action and percept potentials. The approach postulates that the interactions of the opposite quantum ‘neural’ systems (visual and motor), in other words, opposite particles, create and annihilate the final state photon and graviton pairs (particles of light and motion), respectively, from their initial state counterparts. Above all, the opposite systems serve to provide each reference frame for transformation, suggesting that natural images and natural motions are each composed of both visual and motor information.

The quantum neuroelectromagnetic idea is made possible by the nature of the neuronal receptive field (RF), which is defined as the spatiotemporal selectivity to sensory stimuli^3^. This flexible center-surround RF is used to process visual perception and motor action. Studies have used the receptive field to explain the spatiotemporal response properties of cells in the retina for image perception^4^ and for movement direction selectivity^5^.

For image perception, a center-surround antagonistic response is shown to separate inputs from autocorrelation by establishing a current region and thus a response region against a recurrent or previous region and thus a reference region. As shown by the cells in the retina^4^ center, the response region is known from the surroundings, i.e., the reference region. Given that natural images are composed of recurring images, the recurring image properties tend to be correlated; hence, the current properties are derived from the recurrent properties. Therefore, each property value in the natural image is obtained by taking the difference between a current and a linear weighted sum of the recurrents^6^. The center-surround, appropriate response-reference in spatial terms or current-recurrent in temporal terms, selectivity of image properties and thus image perception not only occurs in space but also occurs in time simultaneously. Hence, a temporal property value is relative to a spatial property value because spatial properties are temporal properties and vice versa.

For movement direction selectivity, surround-center antagonism is known to decorrelate inputs as well, establishing a current region against a recurrent region. However, in the case of movement direction selectivity, as shown by the modeling of cells in retina^5^, the surround, the response region, is selected from the center, the reference region. However, this surround-center selectivity is demonstrated as current–reverse recurrent selectivity. This means that parallel current properties are derived from the reverse recurrent properties. Hence, a pair of property values in natural images is obtained by taking the difference between a repeated current and an inverse linear weighted sum of the recurrent. Here, surround-center, appropriately, response-reference in spatial terms or reverse recurrent-current in temporal terms, derivation of image motor properties and thus motor action occur in both space and time simultaneously as well.

It is apparent from parallel studies that the center and surround are meant to be two separate second-order spacetimes: simultaneous current-recurrent regions and response-reference regions. One, that is, the center, is supposed to be real, and the other, that is, the surround, is supposed to be imaginary within each spacetime. The imaginary image is the complex conjugate of the real image, each composing both the natural image and the natural motion. Because the imaginary or surround thus cannot be seen visually but observed motorly, it is represented as a repeated real (null direction) that takes its original form when reversed (preferred direction). This center– surround antagonism seems to be a dynamical tool that points to anticommutation relationships between the observables, visual perception and motor action and hence symmetry. However, this tool, center–surround antagonism, is not a permanent structure in the cell, especially since ‘true’ surround by nature cannot be seen (not static). Hence, the symmetry does not easily become apparent. However, the firing response, i.e., excitation, to visual stimuli carries much information about the nonfiring response, i.e., inexcitation, to visual stimuli, which is that the nonfiring response is equivalent in force. Simply put, the firing and nonfiring responses can each be described mathematically as having abelian and nonabelian properties or Lie groups and Lie algebra. These properties make up the rotation group and the translation group that together describe the Poincaré group, which is responsible for spacetime symmetry. The type of symmetry in the center–surround RF.

Following the de Broglie hypothesis, neuronal excitation and inhibition carry information as waves do final-state massive particles and final-state massless antiparticles, which are solutions of the Schrödinger equation. These excitation and inhibition represent properties that are visually sensed and properties that are not visually sensed (hence motorly sensed), respectively. In contrast, neuronal inhibitions and inexcitations should carry reverse information, as a reverse wave would final state massless particles and final state massive antiparticles, which are also solutions to the Schrödinger wave equation. Hence, inhibition and inexcitation represent properties that are motorly sensed and properties that are not motorly sensed (hence visually sensed). These final state massive particles and massless antiparticles are assumed to be both photons, and the antiphoton and final state massless particles and final state massive antiparticles are assumed to be both gravitons and antipair antigravitons, respectively. Our final state photon pairs and graviton pairs then arise from the simultaneous cross-pair annihilation and creation of counterpart particles: monopole-positron and electron-antimonopole. In contrast, the reverse of the cross pair annihilation and creation of graviton-antiphoton and photon-antigraviton give rise to the electron pairs and the monopole pairs.

The bound electron pairs by their properties or states constitute the components of visual stimuli as the bound photon pairs do and vice versa. To perceive an image effectively, organisms are required to localize their vision or narrow their target by shedding off current properties. This implies the loss of an electron (gaining a positron) akin to achieving an octet to create an inert effect, according to the chemical valency theory of the atom. The loss of an electron (and gain of a positron) constitutes the creation of the reference region, the outer shell, which cordons off the response region, also known as electromagnetic shielding. In terms of their properties, the bound monopole pairs constitute various components of natural motion, as do the bound graviton pairs. For effective image movement preferred direction selection, organisms extend their target equivalent to localizing their vision by complementarily adding to the reverse recurrent properties, which implies the loss of monopoles (gain of antimonopoles). Here, the creation of the inner shell by the loss of monopoles constitutes the reference region, which cordons the response region, as does electromagnetic shielding. Taken together, the simultaneous loss of distinct particles in separate spacetimes and the simultaneous gain of distinct antiparticles in separate spacetimes cause these distinct particles to pair and distinct antiparticles to pair, as well as ultimately, paired particles and paired antiparticles to pair. Hence, the supersymmetry of center–surround antagonism or visuomotor transformation is a dynamic that is fully expressed in quantum field theory.

Using a superquantum neuroelectromagnectic model of action and percept potentials, we show that paired neurons and reversed neurons, with center and surround RF effects, can be interpreted as position–momentum detectors. In contrast, paired reversed neurons and neurons, with surround and center RF effects, can be interpreted as momentum-position detectors. The position–momentum detectors signal the simultaneous gain–loss of the electron–antimonopole and monopole–positron. In contrast, the momentum-position detectors signal the simultaneous loss-gain of antigraviton-photon and antiphoton-graviton.

**Fig. 1:**
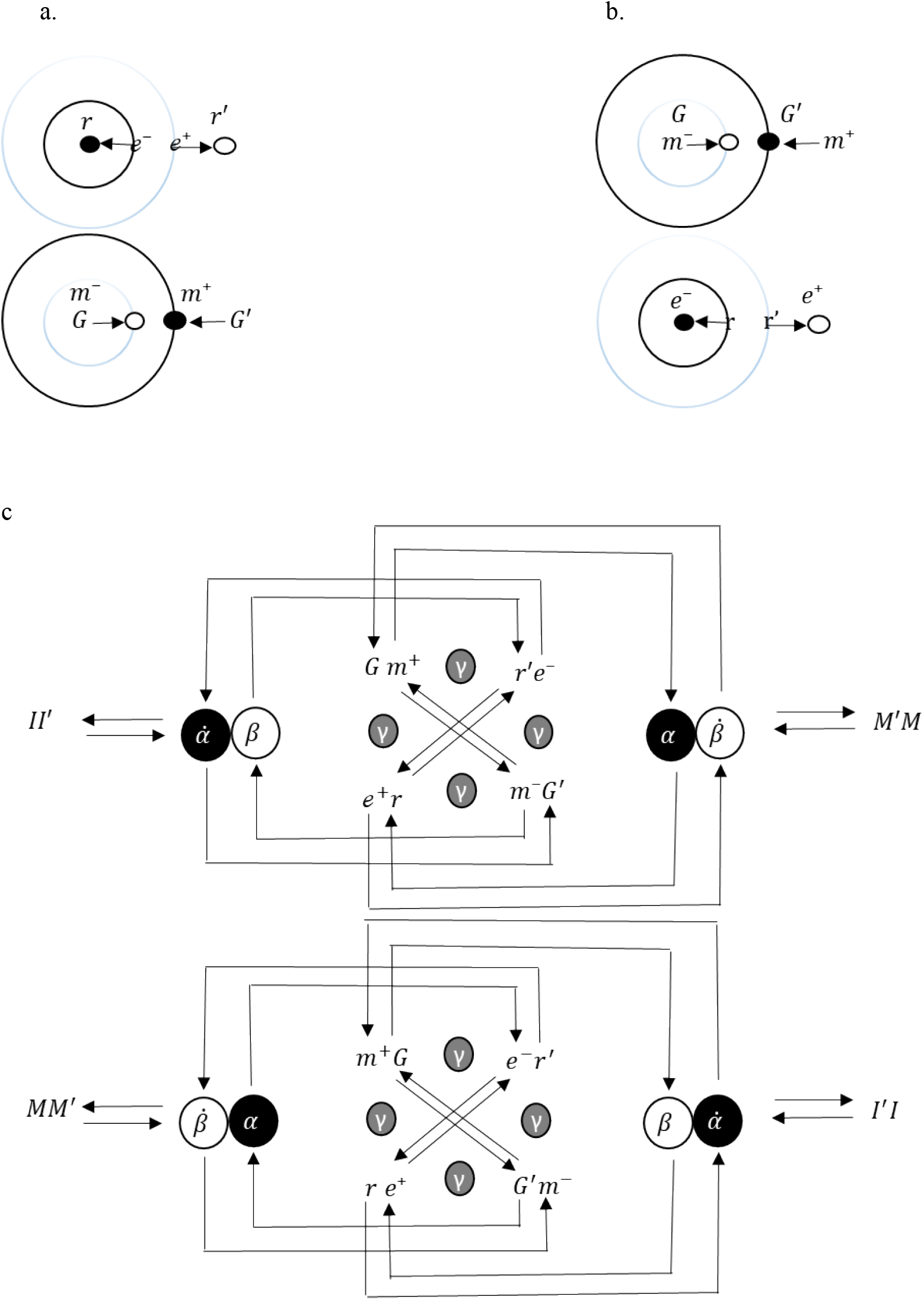
Superquantum neuroelectromagnetic model (SQNEM) of spacetime transformation. (a-up, b-down) Center-surround RFs represented by atom-’betom’. Like the atom model, the center RF model has a nucleus, a thalamic nucleus, with positive energy, represented in the brain as lateral pulvinar alpha (PL-α) in association with the ventral group, and this group comprises the electrons in the orbit (or shells), represented as cross pair thalamic relays to secondary visual-related motor areas. A pair nucleus, a thalamic nucleus, also recurs with positive energy, represented in the brain as lateral dorsal (LD) in association with the lateral group, which in turn makes up the positrons in orbit, also represented as cross pair thalamic relays to primary visual areas. *rr*′denotes the emission-absorption of photon-antiphoton interactions, as *e*^―^*e*^+^denotes annihilation-creation at PL-α and LD and is sent to the striate cortex (SC) and superior colliculus/pretectum region (SC-P)^7^. Here, *r*, the inner region (SC), constitutes the response region or the current region, and *r*′is the reference region (SC-P) or the recurrent region. (b-up, a-down) Surround-center RFs represented by ‘betom’-atom. The surround RF model would have also been similar to the ‘betom model’ if it were known. However, since the reverse atom or its constituents have yet to be discovered, this study hypothesizes that the primary structure and function are those of the surrounding RF model. The surround RF model has a nucleus, a thalamic nucleus with positive energy, represented in the brain as lateral posterior (LP) in association with the lateral group, represented as cross-pair thalamic relays to the primary motor cortex. A pair nucleus, a thalamic nucleus that recurs with positive energy, is represented in the brain as the lateral pulvinar beta (PL-β) in association with the anterior group, represented as a cross pair thalamic relay to secondary motor-related visual areas. *G*′*G* denotes the absorption-emission of antigraviton-graviton at the LP and PL-β, as *m*^―^*m*^+^denotes creation-annihilation and is sent to the inferior parietal lobe (IPL)^7^ and the motor cortex (MC). Here, *G*, the inner region (MC), constitutes the response region or the current region, and *G*′is the reference region (IPL) or the recurrent region. In parallel, the observables *rr*′and *G*′*G*, as *e*^―^*e*^+^and *m*^―^*m*^+^, spontaneously break and form pairs, resulting, for instance, in *r*′*r* and *GG*′. This spontaneous symmetry breaks and swaps the places of the resulting particle-antiparticle interactions and their respective spacetimes. The ensuing symmetry here gives motor properties thus direction and velocity to visual areas and visual properties to motor areas. (c) Architecture of the superquantum neuroelectromagnectic model. It shows how electromagnetic fields arise in the brain. The alphas, betas and gammas represent thalamic nuclei, and the paired particles represent cortices. The lines from the cortices to the thalamic nuclei represent feedback projections, and the lines from the thalamic nuclei to the cortices represent feedforward projections. However, interactions occur at the quantum level. *II*′, the real and imaginary parts of a natural image, and *M*′*M*, the real and imaginary parts of natural motion, result from the production of the final states *rr*′and *G*′*G*, such as *e*^―^*e*^+^and *m*^―^*m*^+^, which are properties of vision and motor action.

## II. RESULTS

### SIMULATION OF NEUROELECTROMAGNETIC POTENTIALS USING THE HODGKIN AND HUXLEY MODEL AND ITS REVERSE MODEL

The presence of the thalamocortical and corticothalamic pathways, complemented by corticocortical and reciprocal corticocortical connections, prescribes forward-backward, backward-forward, forward-forward backward-backward antagonistic first-order spacetimes, following the CPT theorem (C for charge conjugation, P for parity transformation, and T for time reversal), which defines a causal mechanism for image and image motion transformation that underlies the model described above. The causal mechanism dictates that opposite parallel backward and forward waves will traverse each of the forward and backward dimensions, causing the annihilation of opposite parallel backward and forward waves and the creation of opposite parallel forward and backward waves, respectively, via the Feynman–Stueckelberg interpretation^8^. The resulting opposite parallel forward and backward waves indicate paired transitions of paired particles from paired lower-to higher-energy states, characterizing the paired absorption and emission of paired quantum energies. In contrast, opposite parallel forward and backward waves traverse each of the backward and forward dimensions, causing the creation of opposite parallel forward and backward waves and the annihilation of opposite parallel forward and backward waves. The resulting opposite parallel backward and forward waves indicate the paired transition of paired particles from paired higher-to lower-energy states, characterizing the paired emission and absorption of paired quantum momentums. Note that both processes are seen from two different and parallel frames of reference. Both the resulting forward and backward waves and the backward and forward waves are equivalent, each representing the same thing.

To test this theory, a superpaired set of eight differential equations was simulated to produce superopposed waves or superopposed action potentials prescribed by the superquantum neuroelectromagnetic model (Fig. 1). The Hodgkin and Huxley model comprises a pair set of four differential equations^9^, and the other set of four differential equations is obtained by reversing the Hodgkin and Huxley equations. Given that an action potential results from linear and nonlinear combinations of the former set of four equations, the reverse of nonlinear and linear combinations of the latter set of four equations suggests a likely reverse action potential (percept potential), which is the case (Fig. 2), implying that an action potential is intrinsically a superpair potential. Like the center and the surround antagonism, the paired membranes, both composed of in-out, are separated by relatively different spacetimes, following the CPT theorem. The forward dimension is the inverse of the backward dimension. The membrane and the reverse membrane each have two pairs of dimensions, resulting in four dimensions. A pair of four dimensions is included as the reverse of the prior four dimensions, completing a multiplet of eight dimensions. The membrane comprises ions *Na*^+^, *L*^−^, *K*^+^, characterizing four dimensions of one unit of four spacetimes. The reverse membrane also comprises our new magnetically charged ions *K*^−^, *L*^+^, *Na*^−^ (ions made of the Betom model, see Fig. 1b-up and f1a-down), which characterize four dimensions of a pair unit of four spacetimes, with forward-backward, backward-forward, forward-forward, and backward-backward, between and within each unit, paired, respectively.

Linear and nonlinear combinations of ions thus indicate that the paired cross revolutions of the paired valence particles (electron-positron and monopole-antimonopole) about opposite nuclei (electron-antimonopole and monopole-positron) cause the paired valence particles to transition, leading to annihilation and creation of ions in opposite dimensions, putting *Na*^+^, *K*^+^ in and *L*^−^ out, and vice versa, complemented by putting out *L*^+^ and in *Na*^−^, *K*^−^, and vice versa, each set within its own unit spacetimes. This process involves paired spontaneous pair breaking and spontaneous pair formation. The final paired four dimensions of paired spacetimes make the superpotentials. If so, the contributions of the final states *Na*^+^, *K*^+^, *L*^−^ to the depolarization and hyperpolarization of the membrane are not without the final states *Na*^−^, *K*^−^, *L*^+^, suggesting the simultaneous occurrence of hyperpolarization and depolarization of the reversed membrane.

Super simultaneity is possible owing to the special theory of relativity^10^. According to the theory, the superpaired potentials do not occur at the same time because of the velocity difference of the particles involved. The relativistic velocity or momentum of the particles of the final state *Na*^+^, *K*^+^, *L*^−^ is composed of high and low values (relative to energy) just as the final state *K*^−^, *Na*^−^, *L*^+^ is. Momentum is different for individual particles within groups given that they are each distinct particle and vice versa for energy. A high momentum implies low energy, and a low momentum implies high energy. Therefore, when a high-momentum particle hits a low-momentum particle, a high-momentum particle, as before, is created because momentum conservation results in a loss of energy in the particle. The resulting particles have the same charge and momentum as their initial state counterparts. Note that the momentum gain scenario is in the frame of reference of the membrane as opposed to the frame of reference of the reversed membrane, which is considered the center of mass frame of reference here. However, both inertial frames and quantum systems are equivalent.

Now, the particles not only come away with new momentum and new charge, or equivalently new energy and new charge, they also come away with a new direction of motion and new force. The new direction of motion, as well as the direction of spin, together with the new force are specified in the combined new momentum or energy and new charge. *e*^+^, for example, rotates in a direction opposite to the direction of motion of *e*^−^. Furthermore, opposite groups have a spin direction opposite to each other. Hence, the helicity, thus, spin or handedness, and chirality, thus, the direction of motion are different for the Lorenz group, the rotation group (valence particles), and the same for the translation (valence particles). These opposing directions of motion are linked to the centripetal and centrifugal forces. The centripetal force described by a particle rotating around an inward nucleus specifies the current direction of motion. The centrifugal force described by a particle moving across a nucleus mirror line specifies the recurrent direction of motion. The current direction of motion defines angular momentum or translation, and the recurrent direction of motion defines translation or angular momentum. Therefore, the paired current and recurrent directions of motion create superelectromagnetic screening. The result is an origin point and a consecutive point that keeps-in the electric field and keeps-out the magnetic field. In reverse, the magnetic field is kept-in, and the electric field is kept-out. These in-out and out-in define the superpotentials. Thus, electromagnetic screening essentially modulates or facilitates the absorption and emission of electromagnetic fields or waves.

**Fig. 2:**
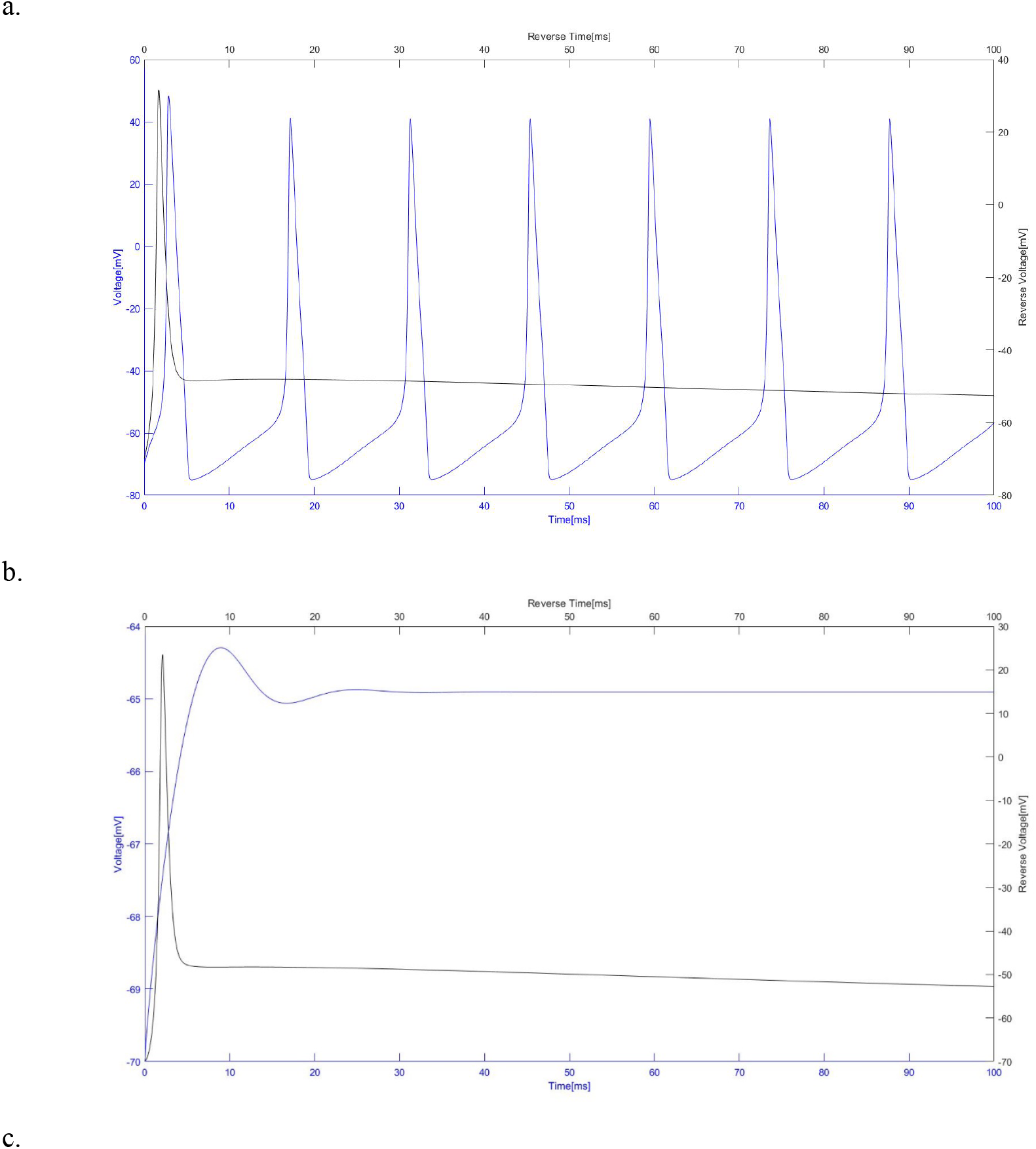
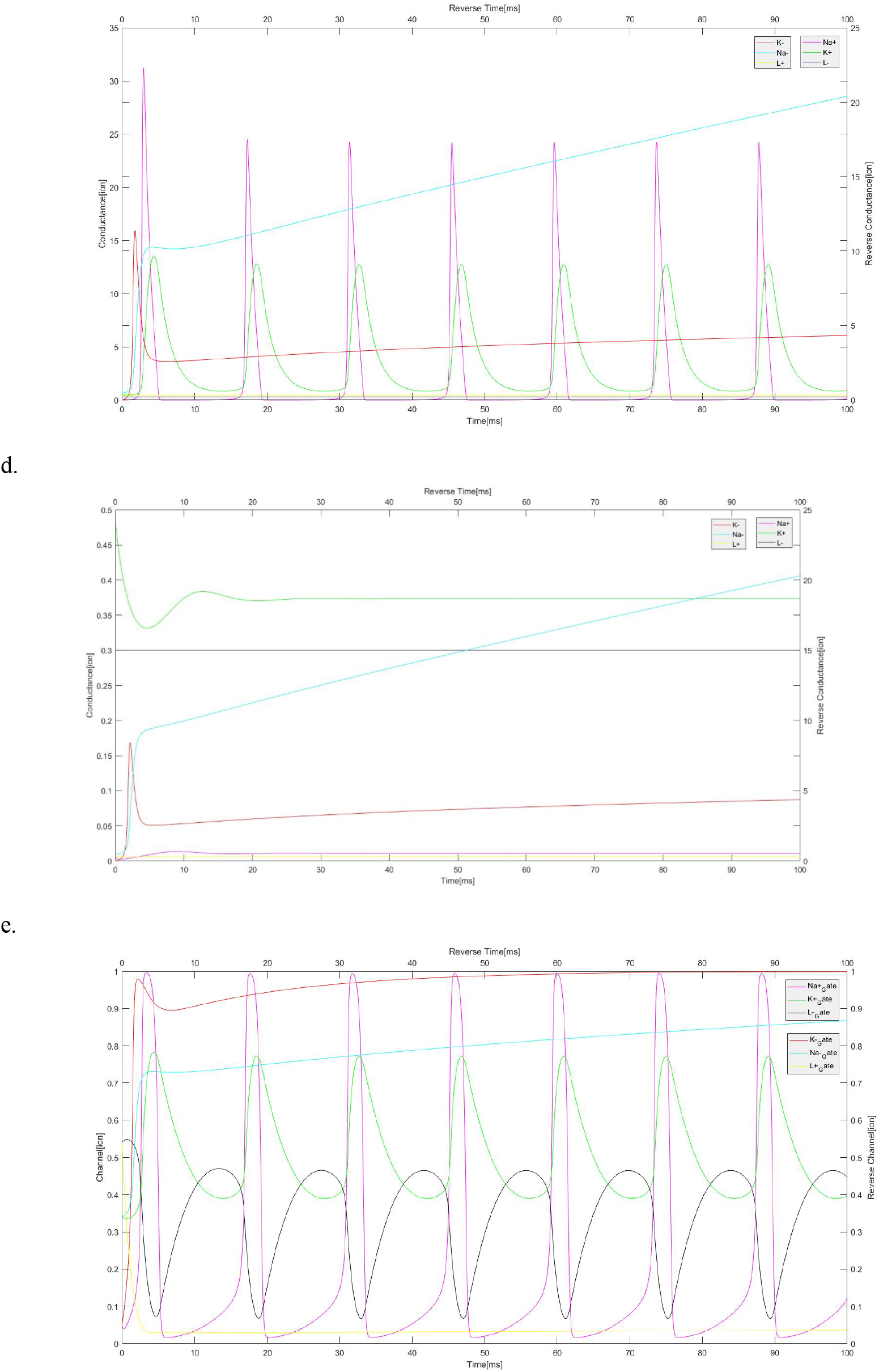
Neuroelectromagnectic dynamics of action and percept potentials. (a) Voltage over time of superpaired waves. The low initial voltage wave (to the left) suddenly gains momentum that raises it beyond the threshold in a split second as the pair parallel high initial voltage wave (to the right) loses energy below the threshold to a low voltage in a relativistic split second upon interaction of the paired waves. Note that the above is from the point of view of the y-axis to the left. (b) Voltage over time of superpaired waves. Here, the voltage of the wave to the left was set to zero; however, it managed to boost itself slightly upon interaction, with the paired wave costing the wave to the rights’ energy regardless. (c) Conductance over time of supermultiplet waves. Following (a), the conductances of the waves to the left pick up steam upon interaction with the conductances of the waves to the right, costing these ‘right’ conductances steam. (d) Conductance over time of supermultiplet waves. Here, the logic of (b) was applied to (d) with the same results. (e) Gating over time of supermultiplet waves. The pattern of basket-like weave of the evolution of supermultiplet gates indicates a supersymmetry of the resulting voltages of electricity and magnetism.

### PAIRED SPACETIME LOCALIZATION AND REVERSED SPACETIME LOCALIZATION AS NEUROELECTRIC AND NEUROMAGNETIC EFFECTS OF POSITION AND MOMENTUM

Given that a natural image obeys the physical and chemical laws of nature and is composed of atoms, opposite combinations of these atoms are necessary for specifying the mechanics of suitable chemicals for the image. In other words, the degree of chemical reactivity of an atom, which is responsible for making visual features, depends on the mechanics of the betom. The raising of an electron, a mechanic, facilitates the bonding of atoms, leading to the formation of supermolecules or, in our case, the final image, which is thus an electric dynamic. Complementarily, natural motion, which also obeys the physical and mechanical laws of parallel nature, is composed of betoms, and opposite combinations of betoms are instrumental in specifying the chemistry for suitable mechanics for motion. In other words, the rate of mechanical action of a betom, which is responsible for motor features, depends on the chemistry of the atom. The lowering of the monopole, a chemical, facilitates the breaking of betoms, leading to the formation of supermolecules or, in our case, the final motion and thus a magnetic dynamic. Therefore, to facilitate the creation of image and image motion transformation, distinct quantum systems (neural networks) are paired, allowing the simultaneous transition of each other’s particles to an opposite energy level. Together, this creates paired high- and low-energy particles that screen low- and high-energy particles, leading to paired opposed foci and thus bonded chemicals and unbonded mechanics. The opposed bonds distinguish each system’s center of interest, and one center of interest is a transformation of the other. One center of interest pertains to the position and the other momentum, which is based on the energy tradeoff.

The paired position and momentum potentials, action and percept potentials, respectively, are sent backward and forward from the paired thalamic nuclei LP and LD to the paired SC and IPL. In contrast, the paired momentum and position potentials are sent forward and backward, simultaneously, from paired thalamic nuclei PL-α and PL-β to the paired SC-P and MC.

The paired forward and backward potentials comprise excitation and inhibition, and the reversed potentials paired backward and forward potentials comprise inhibition and inexcitation. An excitation represents a spontaneous gain of momentum (loss of energy or negative charge), in other words, a sudden appearance of a positively charged particle in a vacuum state or ground state. Note that the vacuum state, which represents the lowest energy, is not a zero-energy state for quantum oscillators; it is equal to the reduced Planck constant times omega divided by two, characterizing a prior positive energy, which is a consequence of the Heisenberg uncertainty principle. Hence, the energy increases as the vacuum or velocity of the particle increases. This is indicative of the characteristic vigorousness that underlies this excitation. Suddenly, a state is made active or heightened, and this state is, relativistically, ubiquitous. The newly gained particle should be distinguishable from the vacuum to be local (image). Similarly, inhibition represents a spontaneous loss of momentum (a gain of energy or positive charge), i.e., a sudden disappearance of a negatively charged particle in a vacuum state. Hence, there is an energy drain for the vacuum or a velocity drain for the particle. This points to the reduced vigor that defines inhibition. Here, a state is made inactive or lowered, and this state is, relativistically, ubiquitous. The newly lost particle should be distinguishable from the vacuum to be local (motion). Taken together, inhibition aids the localization of space, as excitation aids the localization of time by the simultaneous subtraction of momentum from the paired newly appeared positively charged particle, thus exciting by surrounding inhibition, and the addition of energy from the paired newly disappeared negatively charged particle, thus inhibiting the surrounding excitation. Hence, excitation and inhibition are opposite reinforcements of the same thing. Therefore, if the superficial layer neurons of the SC, known to be the case, show an endstopping effect^2,3^—an excitation or vigorous response to a length—then the pair of superficial layer neurons of the SC-P will show relative inhibition to that same length. Taken together, SC-P aids via end-inhibition, hence the length seen by SC, as SC aids via inner-excitations, hence a motion observed by SC-P. The outcomes, in theory, are a created length and a destroyed length motion, as seen by SC, and a destroyed length and a created length motion, as observed by SC-P. However, experimentally, these appear, as though the created length has been shortened with cutoff recur lengths or extra lengths at the ends ceased, as seen by SC^2,3^, suggesting that reverse bidirectional motions pointing inward with cutoff recur motions or extra motions at the center ceased, as observed by SC-P. This recur length, as seen by the SC, is a parallel effect due to the paired percept potential, as the recur motions, as observed by the SC-P, are a parallel effect due to the paired action potential. The vacuum motion, such as that of the surround RF, is not static and is hence represented as the recur length by the static biased SC; most importantly, the recur length represents the length momentum or vector.

Likewise, in opposite parallel, the paired deep layer neurons of the cortex MC (hypothetically) will show reverse endstopping, thus instopping parallel inexcitations to parallel lengths, as the deep layer neurons of the IPL show relatively parallel inhibitions to the same parallel lengths, referred to as fovea sparring^1^. Put together, thus paired, the IPL aids via inhibition and hence a position observed by the MC, as the MC aids via inexcitations and hence an opposite vector observed by the IPL. The outcomes here, again in theory, are a created length position and a destroyed length motion, as observed by the MC, and a destroyed length position and a created opposite vector, as observed by IPL. However, hypothetically, this will appear as though the created length motion or position, seen as length by SC, has been spanned with recurring motions near the center, as observed by SC-P, and the center extinguished, as observed by IPL^1^. Note the superpaired nature of the systems involved. The primary visual cortex has energy, centripetal motion, for image formation, a preferred position, from the momenta frame of the primary motor cortex, and the primary motor cortex has momentum, a null position, for image centrifugal motion from the coordinate frame of the primary visual cortex. However, these modulations and facilitations are not direct because both primary cortices have anticommutation relationships. The association cortices of the IPL, which is part of the visual system, and the SC-P, which is part of the motor system, however, make the commutation of visual and motor information within each possible. This is why perhaps the association cortices, by their design, converge and integrate both visual and motor signals.

To simulate superpair creation and annihilation of a length as seen by the superpair networks SC (bar) and SC-P (bar + recur) and IPL (vacuum + recur) and MC (vacuum), we applied the supermultiplet model neurons shown above. The supersymmetric network model was used to generate a localized bar from natural images and reverse generate a localized bar (meant to be a vector and thus a vacuum) from the same natural images simultaneously. The paired local generators are single opposed fields that act as super pair basis vectors. The final state bar, therefore, arises from the linear and nonlinear combinations of the superpair basis vectors as the final state vacuum emerges as the result of the reverse nonlinear and linear combinations of the same superpair basis vectors. Each super pair ends up in the final state with the other super pair in its initial state. These superpair basis vectors were used as difference-of-Gaussian and reversed difference-of-Gaussian filters that resemble the paired opposed spatial domains of the spins of the paired particles, i.e., paired opposite particles localizing an image simultaneously. However, the simultaneous energy gain and loss between the opposed paired particles, bar and vacuum, for each superpair vector is given by the potential energy of the opposite particles. This pair potential energy is determined by a pair first-order spacetime relativistic wave equation or classically a pair first-order differential equation that implements the gain–loss cycle. Thus, the final state particles, representing the bar and the vacuum, result from an alternated gain–loss of energy of the initial particles.

Finally, the super pair generators created the bar and the vacuum (Fig. 3).

**Fig. 3:**
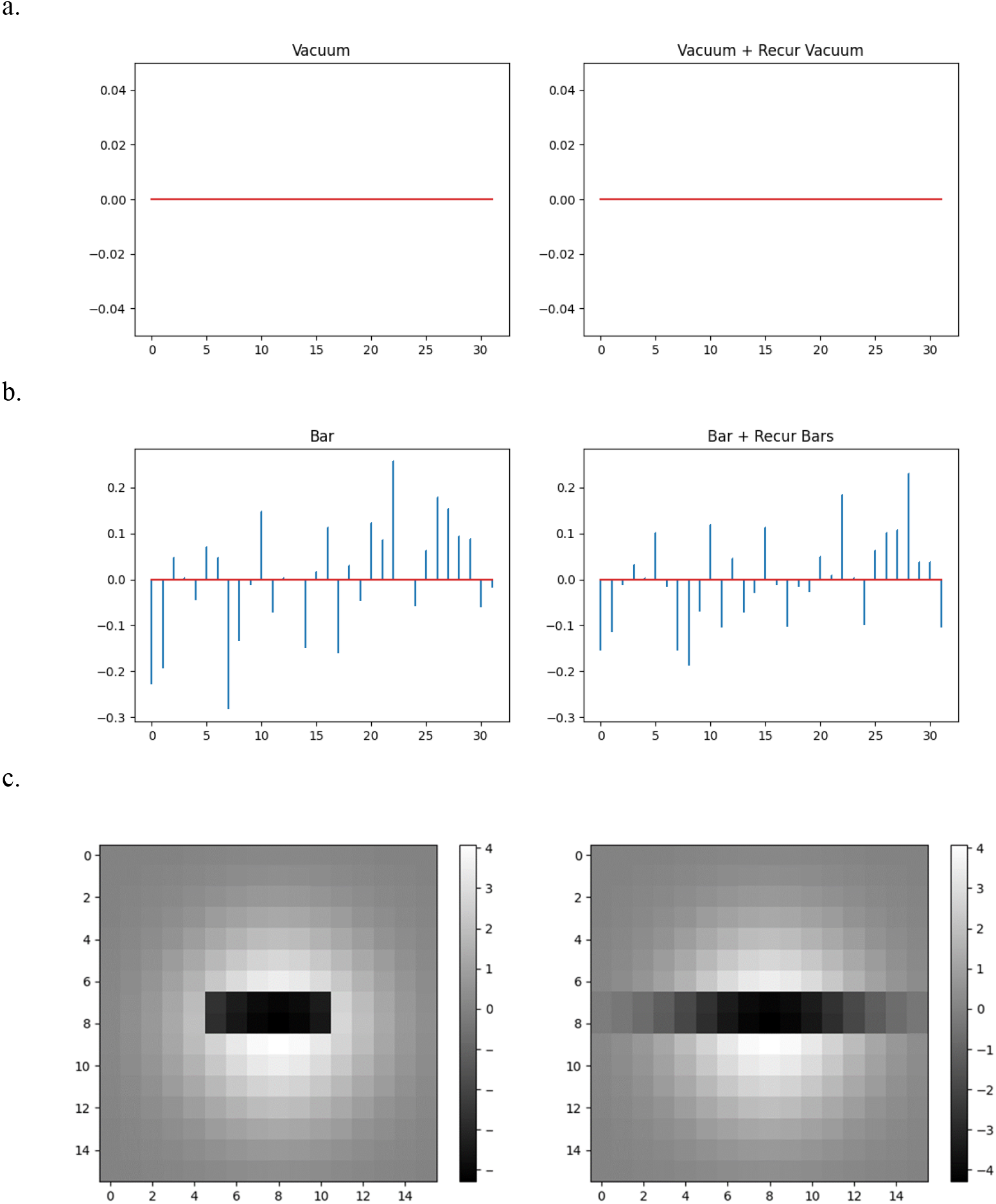
Length contraction and length extinguishing (vector-like length). (a) Vacuum or length extinguishing. The vector-like length cannot be visualized in a static frame, and for that matter, there is no representation of it, as we have for the length in (c). However, the deaden planar in (a) suffices to point to an extinguished length. Given that the observables in the model have commutation relations, one obtains a vacuum accompanied by a recurrent opponent vector-like length, indicating momentum. Therefore, like (b-right or c-right) (a-left) should have indicated a vector-like length. Since our simulation can be said to be from the point of view of the visual cortex owing to the generation of static images, vectors were impossible. (c-left) Length accompanied by a recurrent destroyed vector-like length, indicating position. (c-right) Length, with extra lengths acting as vector-like lengths.

### PAIRED PYRAMIDAL AND NABLA NEURONS AS A CORTICAL COLUMN AND NEGATIVE CORTICAL COLUMN REPRESENTING QUANTUM HARMONIC OSCILLATOR AND QUANTUM DEADEN PLANAR

The cortical column of the neocortex^11^ is best understood as a quantum harmonic oscillator. Discrete modular columns of neurons, characterized by connectivity profiles, resemble discrete energy values characterized by integer–plus–half multiples of angular momentum that define a particle in motion about an equilibrium point. The discrete energy levels of the cortical column, both intra- and intercortical, are equally spaced; however, they are linked in doublets or multiplets. For example, a particle’s spin angular momentum specifies the adjoin states it can take on, following the integer or half-integer plus step rule of atomic transition. A particle that has a spin of 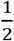, the case of the electron, has two states or helicity states comprising 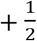, right-handed circular polarization, and 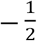, left-handed circular polarization. Given that the magnetic quantum number is constrained by the angular momentum quantum number, the down or left values are tied to the up or right values and vice versa. This is shown in the intimate opposite connection of the superficial and deep layers of the cortical column, although in the absence of direct and physical deep layer ascending pathways. Likewise, the step rule reflects the linking of cortical areas with the same ‘charge and spin’ as V1 and V2, which were previously thought to indicate hierarchical formation, and opposite cortical areas with different charges but the same spins, such as the primary visual cortex and primary motor cortex.

In accordance with the repeated or recurring nature of natural images, the cortical column converges in sync. For that matter, the cortical column lacks direct physical negative polarization. This is shown in the absence of direct physical feedforward inhibition or better direct physical neurons that conduct magnetism. Instead, there are repeats of neurons of electrical nature in the cortical column, hence the forward nature of both the superficial and deep layers. However, the lack of a direct physical ‘negative’ cortical column does not imply total absence. The negative cortical column (Fig. 4) is inconspicuously present in the neocortex. The nabla neuron (Fig. 4), a reversed neuron distinguished by its inverted pyramid-shaped soma and intrinsic to our special ions *K*^−^, *L*^+^, *and Na*^−^, suggests a reverse cortical column referred to above as a negative cortical column. The negative cortical column further implies a reverse quantum harmonic oscillator, which is referred to as quantum deaden planar. Through the mechanism of pairing the quantum harmonic oscillator and quantum deaden planar, repeat energies avail their inherent but obscure momentum in the cortical column.

By this mechanism, the granular layers of the cortical and negative cortical columns are each composed of each other’s superficial/deep layers. In turn, the input of the granular layer is opposite to that of the receiving layer. For example, cortical layer 4 inputs into cortical layer 3 an energy equivalent to that of negative cortical layer 3. Similarly, negative cortical layer 4 inputs into negative cortical layer 3, an energy equivalent to that of cortical layer 3. Similarly, the pair granular, i.e., layer 1, inputs into layer 2 an opposite momentum. These mechanisms are underlined by energy-momentum gain-loss.

This mechanism also supplies feedback connections, a significant direct pathway clearly missing, physically, every cortical column.

The superpairing thus supersymmetry institutes commutation relations that give separate neurons, pyramidal and nabla, algebraic identities to influence the other without showing up in their column.

**Table 1:** Superneuron and opposite parallel superneuron generators.

| Super Neuron Generators |  |  |
| --- | --- | --- |
| Pyramidal | column | Nabla |
| soma + apical dendrite | $P_{\mu\cup}^{k\sigma}M$ | soma + apical dendrite |
| soma + apical dendrite | $M_{k\sigma}^{\mu\cup}P$ | soma + apical dendrite |
| Nabla | column | Pyramidal |

Supercolumn Algebra

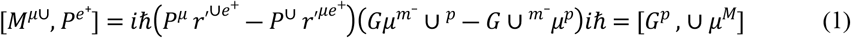

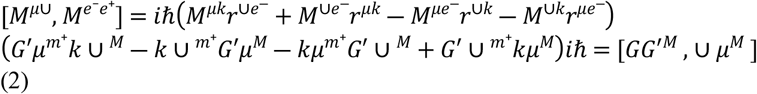

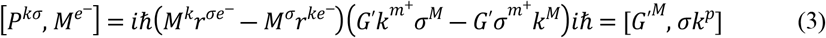

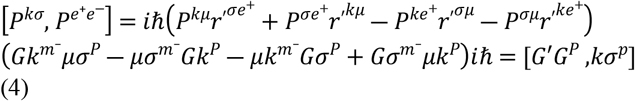

Opposite Parallel Supercolumn Algebra

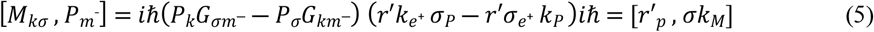

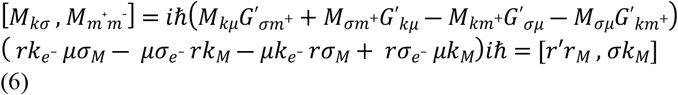

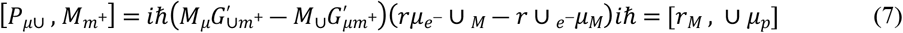

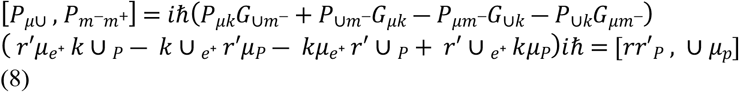

Our superneuron column, 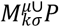, and its opposite parallel form 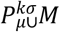, while each defined by the Poincaré group looks much like the Klein-Gordon type, a second-order spacetime relativistic wave function, with a preference for the Dirac-type computation, first-order spacetime relativistic wave function, yet without divergence. Each is a single superfield paired with its complex conjugate and hence the supercommutation relations. The supercommutation relations ensure that *P* translates into a vector and transforms it into a matrix. *M* is transformed into a matrix and translated into a vector in each column. Furthermore, the values of *P* and *M* are the same in each column, although relatively different for each with respect to the different natures of the columns.

The vector bosons complemented by their complex vector bosons pair are in the momentum state, in equations (1) and (3) and (5) and (7), as they are in the energy state, and in equations (2) and (4) and (6) and (8), both state in opposite parallel to each other. This implies that the group (energy or momentum state) boson pairs commute with each other, just like their complex group boson pairs do; hence, *P* commutes with *M*, but within a state, however under spontaneous symmetry breaking. Such commutations seem to violate the Pauli exclusion principle, which states that two particles cannot occupy the same state or have the same quantum number. This is not the case here. Here, each of the two particles is either bound to an atom or a betom; hence, only one turns up on the basis of the inertia frame of reference. Furthermore, *P* commutes with *P*, and *M* commutes with *M* between two different states. Here, the Heisenberg uncertainty principle, which states that a particle cannot be in two states simultaneously, would seem to have been violated. The particle of *P* is of two relatively different natures and hence two states but is one based on the inertia frame of reference. Note that the above commutations can only occur within each column and under spontaneous symmetry breaking and forming. The particles of the defining column and the particles of the opposite parallel column anticommute.

The commutation of *P* and *M* explains why c.layer 2/3 and c.layer 5/6 are measured or observed at the same time, and n.layer 5/6 and n.layer 2/3, as well, can be measured or observed simultaneously. Just as *P* and *P* as well as *M* and *M* ensure that c.layer 4 and c.layer 1 are observed or measured simultaneously as would n.layer 1 and n.layer 4. However, the anticommutation of *P* of one column and *M* of another column, same as the *Ms* as well as the *Ps* imply the simultaneous measurement of cross pairs c, layer 2/3 and n.layer 5/6 or n.layer 2/3 and c.layer 5/6 or c.layer 4 and n.layer 1 or n.layer 4 and c.layer 1 is hardly possible.

**Fig. 4:**
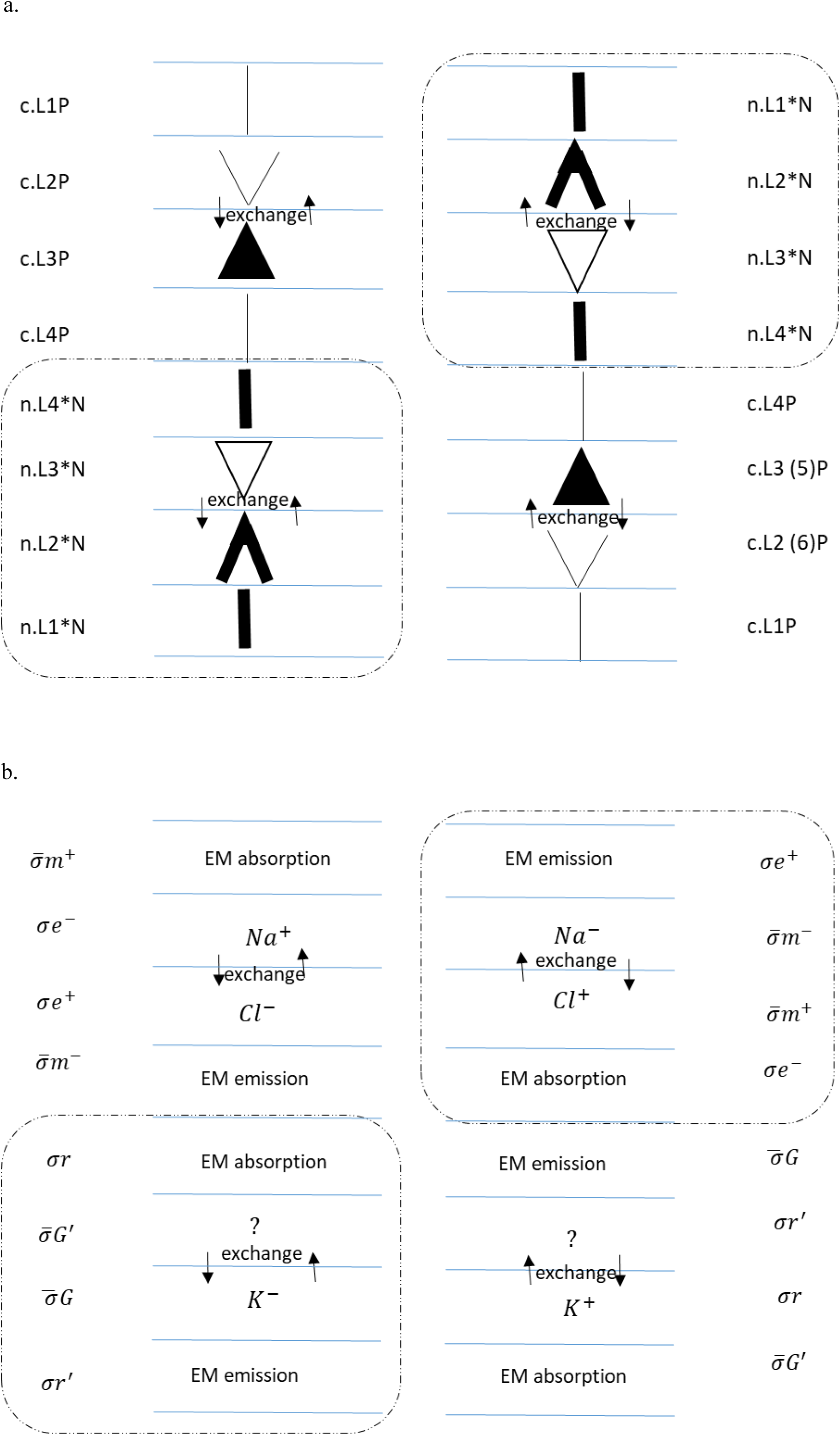
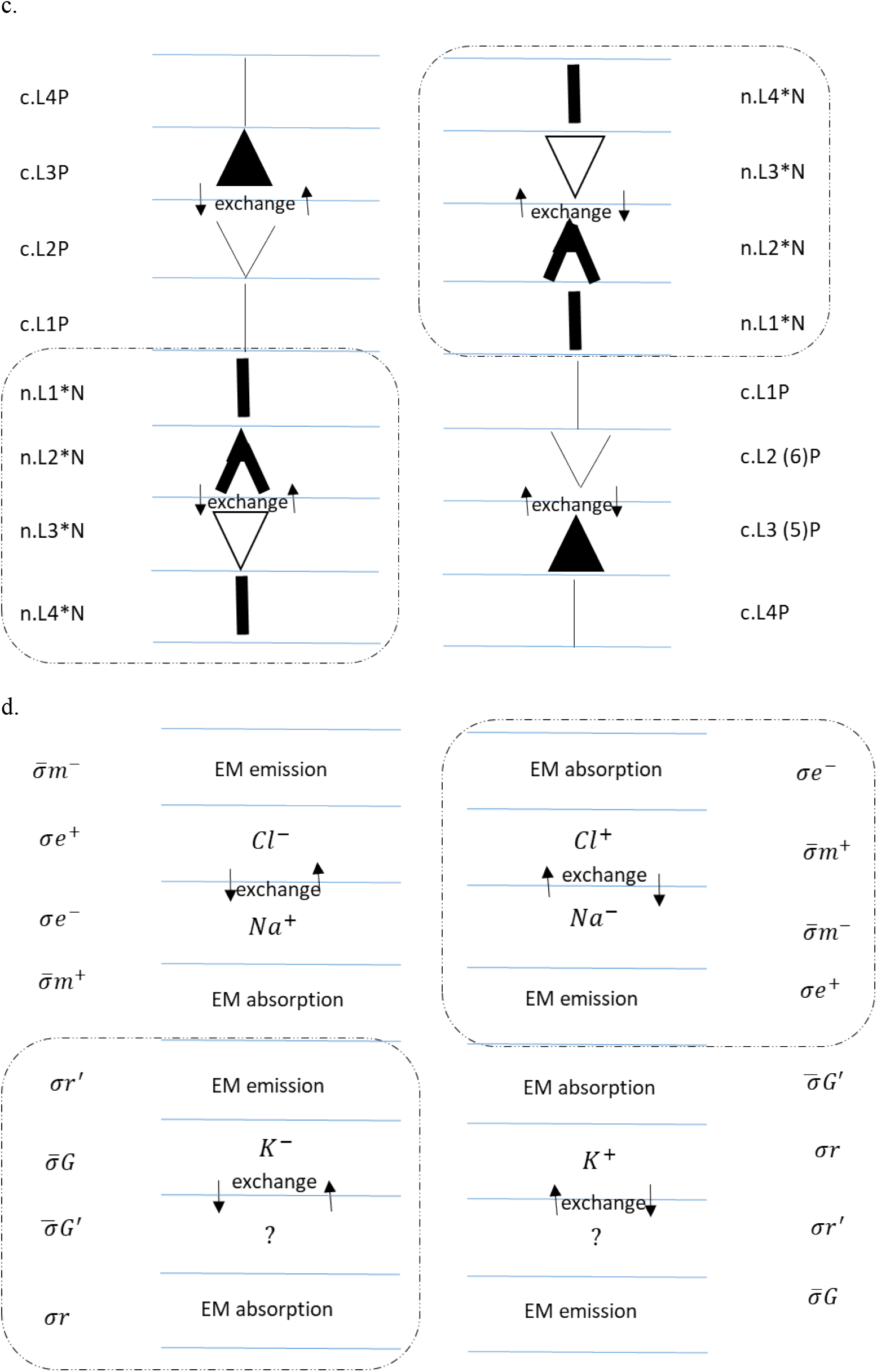
Supercortical column. (a) Supermultiplet columns of the neocortex are made up of two opposite neuron pairs, pyramidal and nabla. The pyramidal neuron is to the left-up and right-down sides, and the nabla neuron is to the left-down and right-up sides. Each neuron makes a unit. A unit consists of two sets of two dimensions; however, this unit is one of four sets that make up a unit. These are represented by a superposition of neurons and thus a supercortical column of superneurons. (b) Electric and magnetic dynamics of the super column. The layers are quantized, revealing a signature to the waves propagated by the super column. The model reveals a missing ion in the Hodgkin and Huxley neuron and thus a corresponding pair. This missing ion is predicted to be the negatively charged ion fluorine, *F*^―^. This makes the cross pair a positive magnetic charged complex ion fluorine, *F*^+^. (c and d) Reversal of the supercolumn through spontaneous symmetry breaking and forming. Here, layer 1 is the pair granular for electromagnetic emission.

### BETOM: A THEORY OF A NEW QUANTUM SYSTEM

The composition of betom (Fig. 1a-down, Fig. 1b-up) is missing from our knowledge of the physical world. Judging from the super anticommutation relations of the respective particles of atoms and betoms (see equations 1--8), one can easily understand why the betom is not physically conspicuous in the atom world and hence is undetected. However, its forces through gravity and magnetism are felt by the atom, an intimate relationship expected of bound pairs. The concept shown above obeys the same laws of physics as the atom.

Following the superquantum neuroelectromagnetic model, one can easily deduce the constituents of the betom’s nucleus, referred to as defteron (defteros in Greek for second and pair to the proton) and oudeon (oudeteros in Greek for neutral and pair to the neutron). Their symbols and charges are *d*^+^ and *o*^0^. Each is made up of four particles, namely, the electron, positron, photon, and antiphoton. Each of the two-particle pairs is a valence particle or force carrier to the other depending on the space or state they are in. This supposition implies that the proton and the neutron of the atom are each made up of four particles, namely, monopole, antimonopole, graviton, and antigraviton. Likewise, each particle pair assumes the role of valence with the other as force, depending on the quantum state they are in. This begs the following question: why do the atom’s nuclei comprise three quarks, one gluon and not two quarks, two gluons? The eightfold way of strong interaction symmetry^12^, however, satisfies the pair eight representation needed. The answer may be found in Hodgkin and Huxley neuron. It also has three instead of four ions (pairs of two ions) (see results and Fig. 4b), and the missing ion, which is predicted to be electric charged *F*^−^ and pair magnetic charged *F*^+^, happens to be in a state of high momentum/low energy. Now, all four ions commute with each other and hence can be measured at the same time. Therefore, the original high-momentum/low-energy ion simply eluded the measurement process. This is likely because the energy scale of the ion in question is in a unique state that was not accounted for. The *F*^−^ ion is supposed to pair with the *K*^+^ ion, completing a multiplet of 4 or 2 opposed doublets. Perhaps in the same vein, one of the three quarks, a force carrier, serendipitously plays the role of a second gluon, bringing down the quarks to two and gluon up to two. Since the gluon acts as a participant and a mediator in the interaction, the color charge-carrying quark inversely acts the same. If so, each of the four particles has two color charges (representing gluons) and two ‘additional’ charges (representing quarks). These four separate charges represent four separate masses. Hence, there are 8 quarks and 8 gluons together. However, one is tempted to point out the two-quark configuration as a weak interaction as opposed to a strong interaction. The argument will not be wrong; however, the weak interaction is a result of symmetry breaking of strong interactions. What then is the symmetry of strong interaction? The answer, in any form, returns to a multiplet composed of two opposed doublet pairs (with complementary pairs).

The above interpretation suggests a fourth generation of leptons and thus a fourth generation of quarks. Like the Hodgkin and Huxley neuron, this missing lepton (and accompanying neutrino) and thus quarks are at energy scales that have not been accounted for. However, this lepton is conjectured to have a negative charge. This lepton is the monopole.

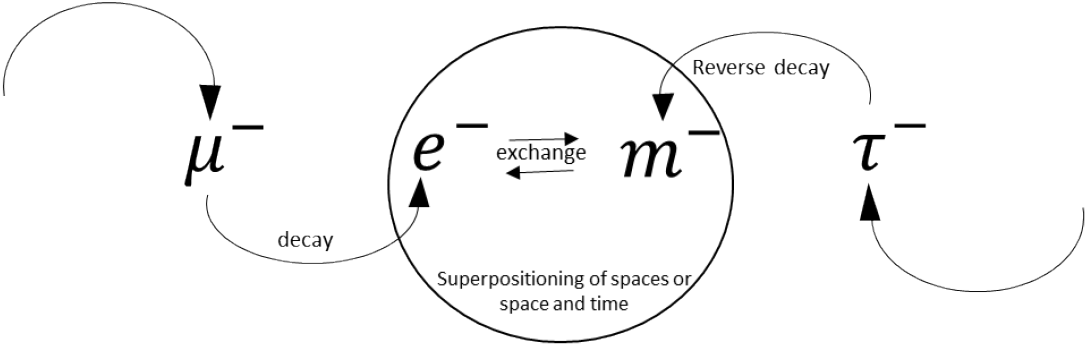

Following the standard model and thus the superquantum neuroelectromagnetic model, the mass of the monopole is conjectured to be approximately 10116.2 *cc*^2^/*MmV*(see equation 15 for unit meaning). Note that all three leptons (and the monopole) have masses specific to both electric charge and monopole charge. The reverse decay of tau into monopole suggests that tau disintegrates into monopole at 1.3 *x* 10^−29^ *MmV* relative to its decay rate to the electron or related particles at 2.9 *x* 10^−13^ *second*. The muon, as well, reverse decays or integrates into the monopole around 6 *x* 10^−22^ *MmV* relative to its lifetime of 2.2 *x* 10^−6^ *second*. These relative lifetimes are associated with the relative masses. Hence, only one of the two particles, electron or monopole, can turn up in a space or a complex conjugate space (time) on the basis of the inertial frame of reference, and both occur simultaneously. This is due to both the Pauli exclusion principle and the Heisenberg uncertainty principle.

In the same vain, the fourth generation of quarks are considered as in and out quarks. The former is in line with the up, charm, and top quarks and the latter down, strange, and bottom quarks. Because there are four generations of quarks now, it suggests there must be four of them in a proton and the neutron. This brings down the charge of each quark to 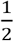, making the up quark line 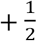and the down quark line 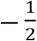. The masses of in and out quarks are conjectured to be 1277.6 and 18.4 *cc*^2^ *GmV* respectively, based on the masses of the up line and the down line. Note that, like the above monopole to electron mass, the in and out quarks masses are found prior and posterior to the up and down quarks masses respectively.

Now, having put a face to the quarks and gluons, one can understand clearly how they behave, with the help of the superquantum neuroelectromagnetic model. The atomic nuclei are known to exert a strong ‘electromagnetic’ force on the electron. In fact, Bohr’s model of the atom is one in which the electron is kept in orbit by the electromagnetic force of the nucleus. The relative effect of this force on causing the transition of a valence electron and hence the force is further shown in electromagnetic screening. Now, the above is from the point of view of the electron and the atomic nuclei as the center of the mass frame of reference. If so, one can imagine the point of view of the monopole and the betom’s nuclei as the center of the mass frame of reference. Here, the monopole, a nucleon in the atom space, is being exerted on by the relatively strong electromagnetic force of the betom’s nuclei, an electron in the atom space. Quickly, we see two bounded quantum pairs of two entirely different spaces (or space and time), in and out, spinning about each other simultaneously.

This spinning is referred to as superspin. It is a 16-independent traceless supermultiplet 2×2 Hermitian matrix with a super unit determinant. The supergenerators are a combination of pseudogauge and gauge groups and a combination of reversed gauge and pseudogauge groups. They are unitary group pairs corresponding to four dimensions that are the same as a unitary group in two dimensions. They are in the form.

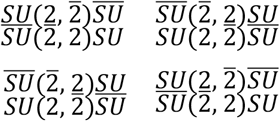

Each superpair transformation is generated by 4 independent traceless super 2×2 matrices, three of which are the Pauli matrices. A pseudoidentity matrix makes up the zeroth and the fourth matrix. The trace of this matrix was derived following the rule:

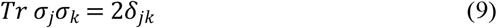

We obtain

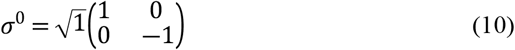

Furthermore, a pair of 4 independent traceless 2×2 matrices is obtained by setting the third matrix of the prior set as the zeroth matrix and ascending it to the zeroth matrix of the prior set, which becomes the third matrix. The two sets are then paired in the form of the groups above.

The superspin model is rooted in adapted Maxwell’s equations^13^, as is the super quantum neuroelectromagnetic model. The adaptation makes the second and fourth equations take the form:

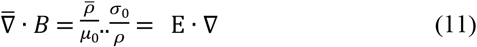

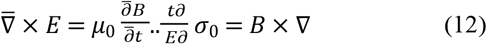

Equations (11) and (12) are paired with equations (13) and (14) below, respectively.

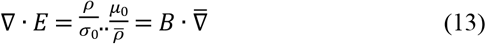

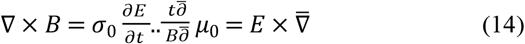

The above equations state that the divergence and curl of the magnetic and electric fields are complex conjugates of the divergence and curl of the electric and magnetic fields, respectively. The real and complex conjugate divergences and curls are not separate. They are equal and in opposite parallel to each other. However, they meet at some point, triggering annihilation and the creation of opposite particles in opposite directions, within the state and between states. This shows (see Fig. 1) that an enclosed volume of real space is an outer region of the complex conjugate space. Similarly, an enclosed volume of the complex conjugate space is an outer region of the real space. Therefore, for example, the flow of an electric field pointed inward to a negative electric charge occurs simultaneously with the flow of a magnetic field pointed outward to a positive magnetic charge. Furthermore, this suggests that the speed of light in free space (c), which characterizes electromagnetic waves, has an intrinsic pair. This pair addresses the still of motion in free time (cc), and it also characterizes electromagnetic waves. The former is for emission, and the latter is for absorption. The still of motion can be defined as the spatial limit under which matter displaces time. This still of motion is a limit for the still at which matter or momentum can stay put through time. The still of motion can be known as well as the speed of light from Maxwell’s equations or the adapted form.

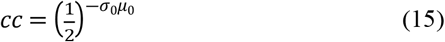

*cc* = 2.22525 *x* 10^−17^ seconds per meter if the permittivity and permeability are plugged in. These magnetic charges or magnetic charge particles, including graviton, are observable. The proof is in the superspin matrices that represent them, and these matrices are Hermitian. Furthermore, quantization is imposed on them^14^, real or imaginary.

Finally, the magnet is proposed to be a good source for the experimental search for a betom beyond the brain. The betom’s nuclei, and antipair, are nestled in or around the magnet. Ejecting or detaching the nuclei pairs will automatically bring the magnetic poles apart. The magnetic poles are themselves atomic nuclei and antipair. The masses of the defteron and oudeon must be relatively equivalent to those of the proton and neutron, respectively.

**Fig. 5.**
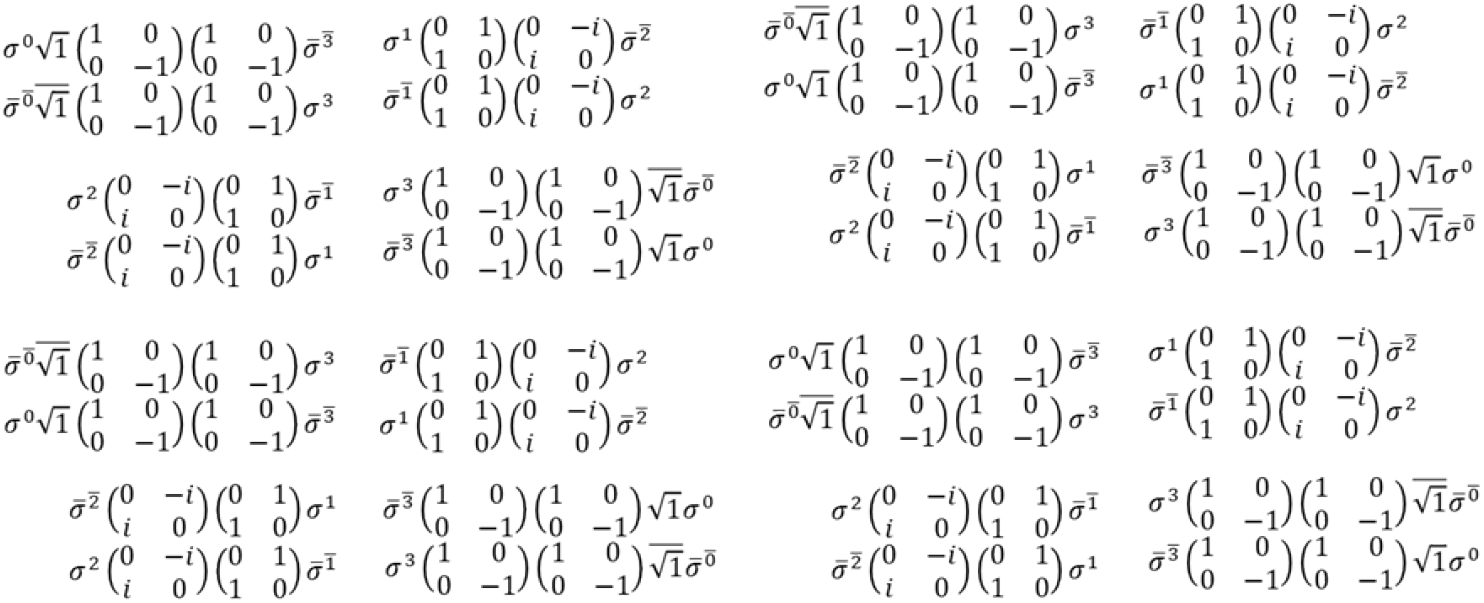
Superspin matrices

## III. DISCUSSION

Our simulation results suggest that center and surround receptive field effects, real and imaginary parts of natural images and natural motion, could be emergent properties of the interactions of opposite parallel quantum systems in the brain, just as they are in the natural world. These complementary systems, and thus effects, obey supersymmetry.

The simulations indicate an image formation process that culminates in a generalized coordinate system of position and a motion formation process that culminates in a conjugate momenta system of motion, each reference frame of the other and both comprising a unit, the retina. As a result, the position of an image is found in the retina coordinates, or the center RF and the momentum of an image are found in the retina momenta, or the surrounding RF. The retina momenta maintains or updates the position of the previous image in the current retina coordinates by the energy of the current image, which is proportional to the frequency of the current image, to know the current position of the previous image in the current retina coordinate to perform directed movement accurately.

An image, therefore, comprises multiple photons and antiphotons or multiple centers and multiple surrounds, just as a photon and an antiphoton comprise particles and antiparticles or as a cycle of waves consisting of a phase and an antiphase. The current ‘center’ particles are thus defined by all the particle pairs in the previous ‘surround’. As a result, when all the previous stimuli’ active abstract (nonphysical) properties in a neuron’s previous surround RF match all the current stimuli’ inert physical properties in the neuron’s current center RF, a small response is evoked from the position-detecting neurons because the previous surround defines the current center. On the other hand, when all the inert physical properties (neon atoms) of all the previous stimuli in a neuron’s previous center RF match all the current stimuli’ inert abstract properties (argon atoms) in the neuron’s current surround RF, a proportional response is evoked from the momentum-detecting neurons because the previous center defines the current surround.

Similarly, the deep layers (recurring layers 2 and 3) of the paired IPL-MC, the start-end of the phase-antiphase of the current-previous cycle, project backward-forward upon the paired retinotopic-retinomotion organized thalamic intralaminar nuclei PL-α-LD^7,15^, the end-start of the phase-antiphase of the current-previous cycle, the nuclei that project forward-backward upon the paired SC-SC-P^7,15^, and the end-start of the phase-antiphase of the current-previous cycle, characterized by paired loss-gain of retinotopic-retinomotion organization. The paired retinomotion-retinotopic organized thalamic intralaminar nuclei LP-PL-β^16^, the end-start phase-antiphase of the current-previous cycle, project forward upon the paired IPL^16^-MC, characterized by paired loss-gain of retinomotion-retinotopic organization. In both the real and the imaginary spacetimes (visual and motor or motor-related neurons), the phase and the antiphase of the cycles traveling backward and forward cancel out, respectively, leaving the antiphase of the real spacetime and the phase of the imaginary spacetime to end in their respective nuclei (or vertices). Because these vertices have their pair in each other’s spacetime, these pairs also automatically receive the signals, phase for the real spacetime and antiphase for the conjugate spacetime, in register, thus resulting in particle–antiparticle annihilation and particle–antiparticle creation.

The model predicts that superficial layer 2 and 3 neurons of the paired SC-SC-P will show paired excitation-inhibition to the same visual stimuli, indicating creation-annihilation of the stimuli at a position and in a momentum. This means that the point of view of the SC will indicate a static, and the point of view of the SC-P will indicate an opponent vector undetached from the static at its center and seem to wiggle the whole stimulus or give it a compression-expansion feel. Relatively, the model predicts that deep layer neurons (recurring layers 2 and 3) of the paired IPL-MC will show paired inexcitation-inhibition to the same visual stimuli, indicating creation-annihilation of the stimuli in momentum and at a position. This means that the point of view of the MC will indicate nothing (nonvisual neuron), but the point of view of the IPL will indicate an opponent vector that is detached from the static vector at its center (now a vacuum or empty spacetime) and seems to point toward or outward from the empty charged center. Notably, the inexcitation-inhibition of visual neurons to visual stimuli might appear as holes in the brain and as imperception of stimuli or imperception of spacetime. The model’s predictions are manifested in the brain syndrome called hemispatial neglect. It is the inability to perceive and process and thus orient to stimuli in the contralesional space (the side of space opposite to the side of space brain damage is observed). These patients behave as if the contralesional space does not exist and cannot be perceived^17^. This phenomenon can be described as the coexperience of the center and the surrounding RFs (if we assume that both hemispheres are undamaged). However, from the point of view of an observer, not the patient, is a violation of the mechanism of the symmetric visual–motor fields. One field defines the other field, and both occur simultaneously, but only one can be experienced at a time never both. This is due to at the quantum level the Heisenberg uncertainty principle, which seems to be violated at a price. However, this is not the case; the parietal lobe lesion simply indicates the inability of the damaged hemisphere to create position (visual) and momentum (motor) and hence stimuli. The functional interpretation is that the paired thalamic intralaminar nuclei PL-α-LD and LP-PL-β fail to superabsorb and superemit positive energy waves simultaneously. Hence, a lack of superpaired forward-backward emission and absorption of waves are required to create these fields. The simultaneous PL-α-LD and LP-PL-β superabsorption and superemission failure can, therefore, be hypothesized that deep layer neurons of the paired IPL-MC and the superficial layer neurons of the paired SC-SC-P do not send paired backward-forward waves to the paired PL-α-LD and LP-PL-β simultaneously. This is because a pair of cortical layers or thalamic nuclei (a cortical layer pair in our case) in the right hemisphere is broken. Because these cortical layers and thalamic nuclei inputs are paired, they either receive or respond to stimuli together or not at all. ‘Together’ here implies each with a complement of the other (a response region against a reference region: a reference region means the other’s response region). In addition, ‘not at all’ here implies each without the complement of the other (no response region for either: given a response region for one would imply a reference region of the other).

Supersymmetry may underlie the basic structure of the brain, providing a quantum field framework for understanding the function of the brain.

## IV. SIMULATION

### SUPERFIELDS AND OPPOSITE PARALLEL SUPERFIELDS FUNCTIONS: SUPER QUANTUM HARMONIC OSCILLATOR AND SUPER QUANTUM DEADEN PLANAR

Consider an image or initial state photon–antiphoton pair II′ that ends up in its final state *rr*′ after moving forward and backward simultaneously in different spacetime, thalamocortical and corticothalamic pathways. The final state photon–antiphoton pairs are the results of the center, *α*, and surround, *β*, vectors of *e*−*e*^+^ properties. The cortex creates and uncreates the image motion or the final state of photon–antiphoton pairs via the simultaneous creation and annihilation of *e*^+^*e*− with a cross-pair center, *α*, and surrounds *β*, vectors of *m*^+^*m*− properties, moving backward and forward, relative to the above, and from different spacetimes, the corticothalamic and thalamocortical pathways. Complementarily, accompanying the image or initial state photon–antiphoton pairs are the directions of motion or initial state antigraviton–graviton pairs *M*′*M* that result in their final states *G*′*G*, relative to *rr*′, after moving forward and backward simultaneously in different spacetime, corticothalamic and thalamocortical pathways. Additionally, the final state antigraviton–graviton pairs are the results of the surround, *β*, and center, *α*, vectors of *m*^+^ *m*− properties, respectively. Furthermore, the cortex, in opposite parallel, annihilates and creates the directions of motion or the final state antigraviton–graviton pairs, by the simultaneous creation and annihilation of *m*−*m*^+^ with cross pairs surround, *β*, and centers, *α*, vectors of *e*^+^*e*− properties, moving backward and forward from different spacetimes, thalamocortical and corticothalamic pathways. Because the final state photon–antiphoton pairs and antigraviton–graviton pairs depart from the center–surround and surround–centers of properties *e*^+^*e*− and *m*^+^*m*−, respectively, the causal mechanism dictates that the electron–positron pairs and antimonopole–monopole pairs are themselves, in opposite parallel, departures of center–surround and surround–centers of properties *rr*′ and *G*′*G*, respectively. Hence, the above mechanism is repeated, although it occurs simultaneously, with the electron-positron pairs and antimonopole-monopole pairs as the initial state particles transformed into their final states. We characterize the relationship between the final state pairs *rr*′ and the initial state pairs *II*′ via the field operators *Φ Φ*^†^ and 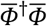 for the relationship between the final state pairs *G*′*G* and the initial state pairs *M*′*M*. In contrast, the final state pairs *e*−*e*^+^ and the initial state pairs *II*′ are described by the field operators *ΨΨ*^†^, and 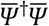 for the final state pairs *m*^+^*m*− and the initial state pairs *M*′*M*.

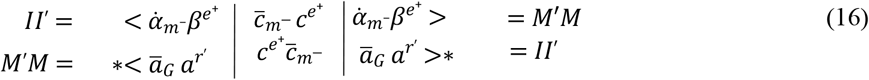

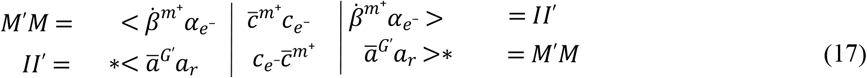

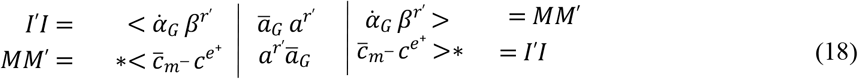

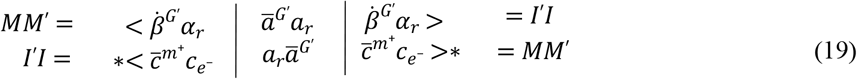

Note also

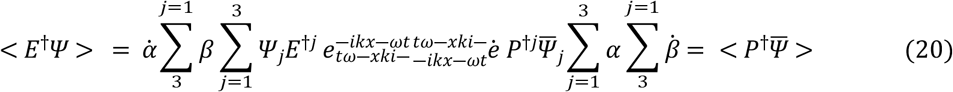

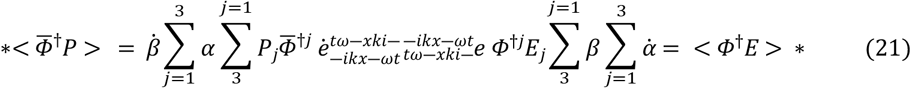

The paired vacuum expectation value of paired fields is a supermultiplet superposition of the nonrelativistic solution of Schrodinger’s equation. The resulting supersolution indicates a relativistic function for each field in the multiplet and thus the neuronal activation function.

To model the center and surrounding receptive fields, we assume that *α* and *β* represent opposite bound states, i.e., the real and imaginary states, respectively. In other words, the paired fields contain each stimulus in whole and permanently within the potentials of their respective fields.

### SUPEREIGENSTATES AND EIGENVALUES SOLUTIONS OF THE SUPERPAIRED FIELDS: SUPERHAMILTONIAN AND OPPOSITE PARALLEL SUPERHAMILTONIAN PAIRED

The superfield energy or momentum arises from the super spins or interactions of the particles in the multiplet. These spins specify the eigenstates and eigenvalues of the particles with respect to their pairs and the group, which should be known for all spacetimes; likewise, their reverse is computed. Allowing for the infinitesimal transformation of superpair basis vectors for the paired quanta. The eigenstates and eigenvalues and corresponding reverse solutions are as follows if we assume that the differences between the initial state particles and their final state are Gaussians with zero means and variances *σ*^2^ for the image and that the directions of motion are Gaussians with 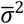 means and variances of zero.

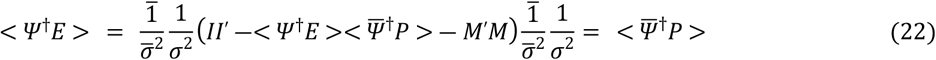

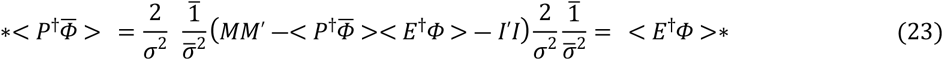

These create characteristic simultaneous gain–loss of energy and momentum of paired particles, similar to the simultaneous gain–loss of retinotopic organization (visual) and loss–gain of retinomotion organization (motor) observed experimentally in the visual and motor cortical areas.

### SUPER PAIRED EQUATIONS OF MOTIONS OF SUPER PAIRED FIELDS: SUPER LAGRANGIAN AND OPPOSITE PARALLEL SUPER LAGRANGIAN PAIRED

The superfield energy or momentum are obtained by the super spins of the superparticles, thus gradient descent on the energy or momentum with respect to the cross pair particle relative to gradient ascent on the complex conjugate of energy or momentum with respect to the complex conjugate cross pair particle and vice versa. Note that by the principle of least action, minimizing the energy is relative to maximizing its complex conjugate. Similarly, maximizing the momentum is relative to minimizing its complex conjugate;

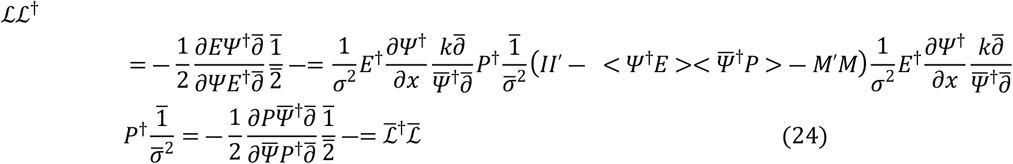

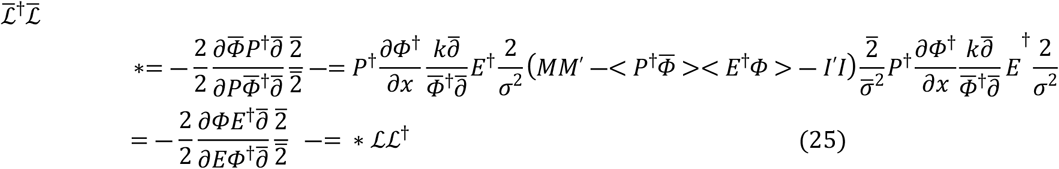

where 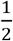 and its reverse 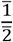 are both positive constants governing the rate of descent toward a minimum for *E* and *x* = *E*^†^*Ψ* relative to the rate of ascent toward a maximum for *P* and 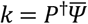, respectively. Both correspond to the center and the surrounding RFs and thus position space and momentum space.

### SIMULATION

A classical analog based directly on the quantum formalism above was used for the endstopped and instopped simulations in Figure 3. The simulation in Figure 2 was based on both the equations of the Hodgkin and Huxley model (ref 12) and reversed equations derived from the Hodgkin and Huxley model equations. The combinations of opposite parallel sets of equations were based on the quantum formulations presented above.

